# Hindbrain neuropore tissue geometry determines asymmetric cell-mediated closure dynamics

**DOI:** 10.1101/2020.11.02.364513

**Authors:** Eirini Maniou, Michael F Staddon, Abigail Marshall, Nicholas DE Greene, Andrew J Copp, Shiladitya Banerjee, Gabriel L Galea

## Abstract

Gap closure is a common morphogenetic process. In mammals, failure to close the embryonic hindbrain neuropore (HNP) gap causes fatal anencephaly. We observed that surface ectoderm cells surrounding the mouse HNP assemble high-tension actomyosin purse-strings at their leading edge and establish the initial contacts across the embryonic midline. The HNP gap closes asymmetrically, faster from its rostral than caudal extreme, while maintaining an elongated aspect ratio. Cell-based physical modelling identifies two closure mechanisms sufficient to describe tissue-level HNP closure dynamics; purse-string contraction and directional cell crawling. Combining both closure mechanisms hastens gap closure and produces a constant rate of gap shortening. Purse-string contraction reduces, whereas crawling increases gap aspect ratio, and their combination maintains it. Closure rate asymmetry can be explained by embryo tissue geometry, namely a narrower rostral gap apex. At the cellular level, our model predicts highly directional cell migration with a constant rate of cells leaving the HNP rim. These behaviours are reproducibly live-imaged in mouse embryos. Thus, mammalian embryos coordinate cellular and tissue-level mechanics to achieve this critical gap closure event.

## Introduction

Closure of embryonic tissue gaps is a common morphogenetic process critical to the formation of structures including the eyelids (1), palate (2), body wall (3) and neural tube (NT) (4). The process of NT closure has long served as a paradigm of morphogenesis and remains clinically relevant today. Failure of NT closure causes severe neurodevelopmental defects in around 0.1% of human pregnancies globally (5, 6). Despite their clinical importance, the cellular force-generating mechanisms which deform embryonic tissues to close the NT are poorly understood.

NT closure starts with V-shaped bending of the flat neural plate at the hindbrain-cervical boundary, elevating lateral neural folds which meet at the dorsal midline (4). The point at which the neural fold tips first meet is called Closure 1. Without Closure 1, the hindbrain and spinal NT remain open, producing craniorachischisis (4). Absence of Closure 1 formation is characteristic of homozygous mutations in core planar cell polarity components such as *Vangl2* (7–10). Soon after Closure 1 forms, a distinct elevation and midline apposition process at the midbrain-forebrain boundary establishes Closure 2 in mice. The resulting gap of open NT between Closure 1 caudally and Closure 2 rostrally is called the hindbrain neuropore (HNP) (4). Midline fusion points, referred to as “zippering” points, form at each of these closure sites and progress towards each other, completing closure when they meet. Note that in the context of NT biology the term zippering is used to denote tissue-level propagation of closure from a pre-existing contact point, as distinct from a “buttoning” process whereby multiple midline contacts form simultaneously (11).

Failure of the HNP to close produces the fatal defect exencephaly/anencephaly. This pathological endpoint is observed in a large number of transgenic and teratogenic mouse models, likely reflecting a multitude of underlying genetic and molecular processes (12, 13). Similarly, at the cellular level, there remain many unanswered questions about how HNP closure is achieved, including the relative importance of individual genetic mutations identified in anencephalic human foetuses. Unanswered fundamental questions include the relative importance of individual cell behaviours to achieving closure and how these are collectively integrated at the tissue scale. Closure rate dynamics are hypothesized to modulate anencephaly risk by determining whether closure completes before a developmental “deadline,” after which tissue-level changes preclude closure (14).

Our recent studies of NT closure in the presumptive spinal region establish that it is a biomechanical event involving both the neuroepithelium and non-neural surface ectoderm (15, 16). The spinal NT surface ectoderm cells assemble high-tension actomyosin cables which border the region yet to close (15, 17), but whether similar cables are present in the HNP is unknown. Genetic and live-imaging evidence also implicates the surface ectoderm in HNP closure, as deletion of surface ectoderm genes such as Par1/2 (18) and Grhl3 (19, 20) produces exencephaly. More directly, surface ectoderm cells produce delicate cellular ruffles which extend across the embryonic midline to meet their contralateral equivalents (21–23). These projections are proposed to mediate midline fusion (24). Detailed analysis of equivalent protrusions in the spinal NT found they are genetically controlled by actomyosin cytoskeletal Rho-GTPase regulators (25).

Genetic deletion or pharmacological inhibition of actomyosin and its regulators commonly produces exencephaly in mice (26–29). Actomyosin contractility, turnover and cytoskeletal assembly determine tissue tension in neurulation-stage vertebrate embryos (15, 30). This contractile network is interlinked between multiple cells in epithelia through direct anchoring to adherens junctions. Supra-cellular actomyosin sheets or cables allow cells to act collectively in deforming their tissues (31, 32). In the studies presented here, we identify and biomechanically characterise surface ectoderm actomyosin purse-strings which contribute to HNP closure, and test their contributions using cell-based physical modelling corroborated by mouse embryo live imaging. We find that a combination of purse-string contractility and directional cell crawling ensure timely HNP closure, but tissue geometry constrains rostrally-directed closure, producing a rate asymmetry in cell-mediated closure.

## Results

### Surface ectoderm purse-strings encircle the closing mammalian HNP

Actomyosin purse-strings demarcate the HNP rim throughout the period of closure (Fig. 1A-C). They line the neural fold tips at the boundary between the surface ectoderm and neuroepithelium (Fig. 1A). At early stages of closure, each purse-string cable extends over 600 μm rostro-caudally (Fig. 1B). High resolution imaging demonstrates cable co-localisation with the surface ectoderm marker E-cadherin, thereby demonstrating their presence in surface ectoderm cells (Fig. 1D). F-actin rich membrane ruffles are also visible along the length of the HNP (Fig. 1D), extending from the cable-producing cell borders. These ruffles are reminiscent of the membrane protrusions characteristic of migrating cells (33, 34).

**Figure 1:**
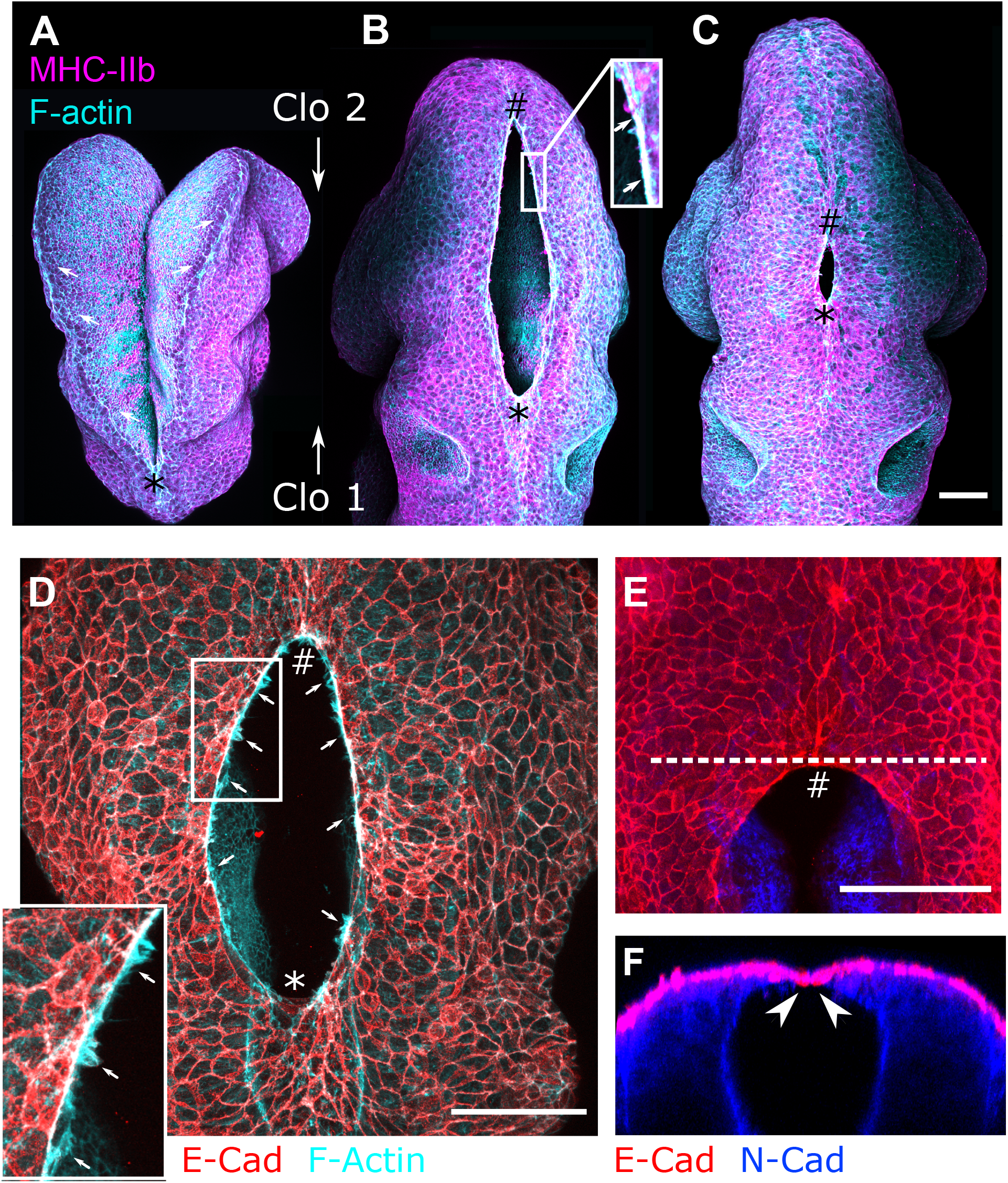
Surface ectoderm purse-strings encircle the mammalian HNP. Representative whole-mount images of the developing cranial region in the mouse embryo. Throughout, zippering progression from Closure 2 (# rostral) is at the top and progression from Closure 1 (* caudal) is at the bottom of the image. **A-C.** Dorsal views of representative embryos before **(A)** and after **(B, C)** HNP formation. **A.** Pre-HNP embryo (E8.0, 8 somites) illustrating progression of zippering from Closure 1 (*). Actomyosin cable-like enrichments (arrows) are detected along the open neural folds. **B.** Embryo with an early HNP (E8.5, 15 somites). The actomyosin cables become purse string-like. Inset: Actin and myosin colocalisation in the encircling cable (arrows). **C**. Embryo approaching completion of HNP closure (E9.0, 17 somites). Directions of zippering from Closures 1 and 2 indicated by * and # respectively. **D.** Co-localisation of cable F-actin with the surface ectoderm marker E-cadherin. Inset: membrane F-actin-rich ruffles (arrows) which appear to extend from the actomyosin cables. **E-F.** Surface ectoderm cells make the first point of contact at the zippering point. **E**. Representative dorsal view of the Closure 2 zippering point with the neuroepithelium stained with N-cadherin. **F.** Optical reslice along the dashed line shown in (E); arrowheads indicate the first point of contact, which is between E-Cad positive surface ectoderm cells.

**Table 1:** Final model parameters.

| Parameter | Symbol | Value |
| --- | --- | --- |
| Contractility | $\Gamma$ | 0.04 |
| Preferred perimeter | $P_0$ | 3.5 |
| Purse-string tension | $\Lambda_{ps}$ | 0.143 |
| Cell crawl speed | $v_0$ | 0.01 |
| Drag coefficient | $\mu$ | 5 |
| T1 length | $L_{T1}$ | 0.1 |
| Time step | $\Delta t$ | 0.2 |

E-cadherin positive junctions initiate contacts across the embryonic midline at the HNP zippering points (Fig. 1E, F). These sit on top of a fibronectin-containing ECM which extends to the surface ectoderm leading-edge, providing a potential adhesion substrate (Video S1). Evidence of ECM anchorage, purse-string assembly and previously-reported midline partner intercalation (23) suggest distinct potential cellular mechanisms of HNP closure.

### HNP closure progresses faster caudally then rostrally

We next documented tissue-level dynamics of HNP closure using a series of fixed embryos and live-imaging in whole embryo culture. During HNP closure, both the length and width of the open region decrease with advancing somite stage (“developmental time”, Fig. 2A-C). The resulting shape of the HNP is initially highly elongated, with a length/width aspect ratio exceeding 3. This aspect ratio decreases slightly but maintains an elongated shape throughout the closure period (aspect ratio > 2, Fig. 2D). Over this time, the zippering point originally from Closure 1 moves rostrally, whereas the equivalent point from Closure 2 moves caudally. Their relative contribution to closure was inferred by calculating the distance between each zippering point and the developing otic pits, used as normalization landmarks in fixed embryos (Fig. 2E). HNP zippering from Closure 2 progresses ~400 μm within 12 hours (2 hours per somite, closure over somite stages 12-17) compared with ~200 μm from Closure 1 on average (Fig. 2E-F).

**Figure 2:**
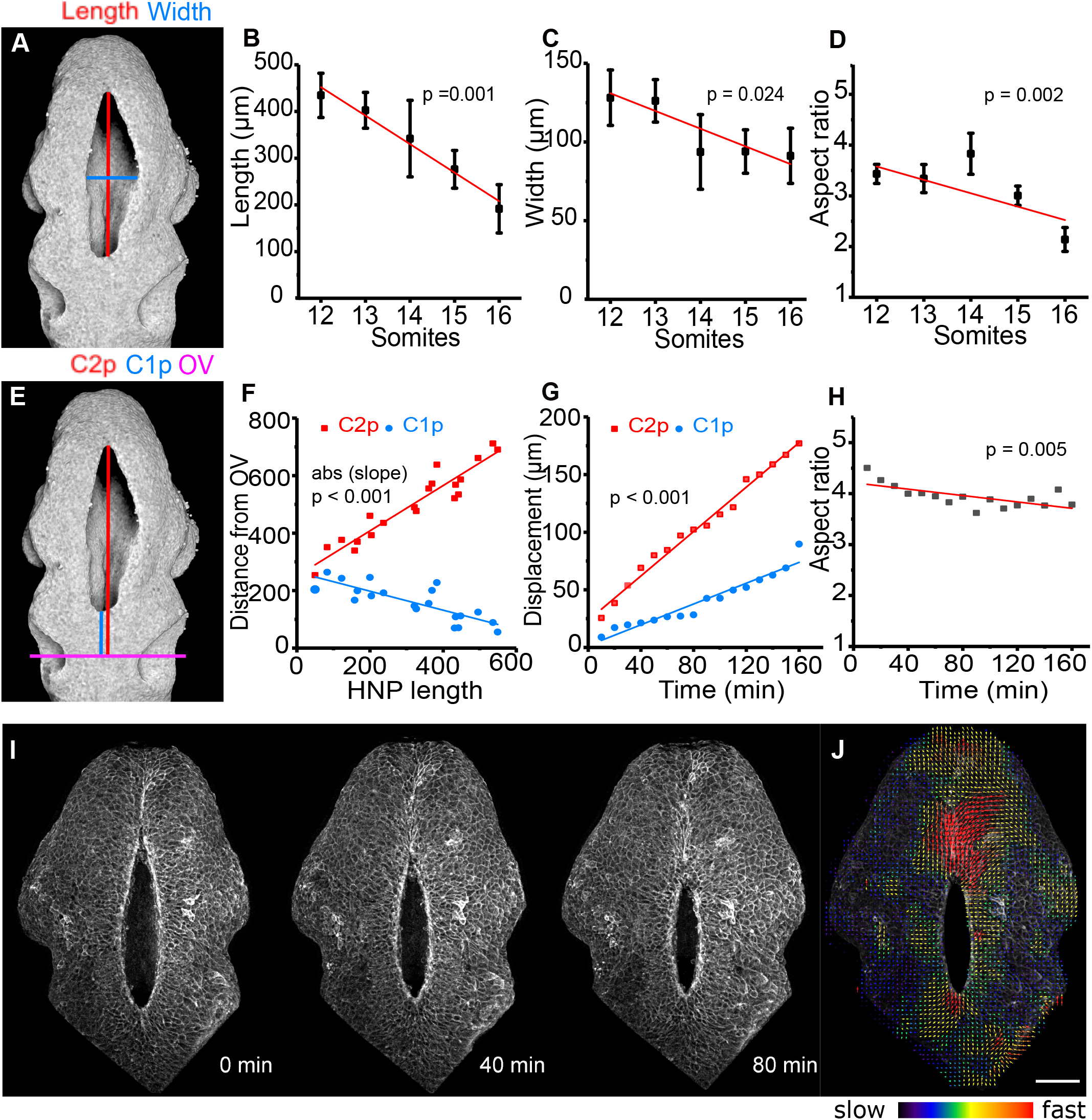
HNP closure progresses faster in the caudal than in the rostral direction, while HNP shape remains elliptical. **A.** 3D reconstruction of a representative early HNP reflection image. The red and blue lines indicate HNP length and width respectively. **B-D:** Quantification of HNP length **(B)**, width **(C)** and aspect ratio **(D)** in fixed embryos at the indicated somite stages (n = 40). The HNP shortens and narrows as it closes, but it maintains an aspect ratio greater than 2. In all cases, the slope is significantly different from 0 (p<0.05, ANOVA) indicating reduction with advancing somite stage. **E.** Same image as in A. The magenta line shows the mid otic vesicle (OV) level. The red and blue lines show the distance of closure 2 progression (C2p) and closure 1 progression (C1p) from mid-OV respectively. **F.** Quantification of the distances shown in E against HNP length in fixed embryos (n= 20). The absolute slopes of the regression lines are significantly different from one another. P< 0.001, F-test. **G.** Quantification of C2p and C1p displacement over time in a live-imaged embryo (shown in I). The slopes of the regression lines are significantly different from one another. P< 0.001, F-test. **H.** Quantification of HNP aspect ratio over time in the same live-imaged embryo (I). The HNP maintains a highly elliptical shape throughout closure. **I.** Snapshots of a live-imaged mTmG mouse embryo at the time points indicated (~15 somites at first frame). **J.** Particle image velocimetry (PIV) illustrating increased cell speed at the HNP apices. Scale bar: 100 μm.

These two features of HNP closure dynamics, namely faster rostral to caudal progression of closure while maintaining an elongated aspect ratio, are also observed during live-imaging (Fig. 2G-I and Video S2). Particle image velocimetry (PIV) analysis suggests wide-ranging cell displacement around the HNP rim, with the highest velocities at the zippering points (Fig. 2J).

Asymmetry in the rate of closure means progression from Closure 2 is responsible for forming a larger proportion of the HNP-derived roof plate and raises the possibility that the two closure points act independently of each other. We next tested closure point inter-dependence directly using a transgenic model which lacks Closure 1.

### Progression of HNP closure from Closure 2 is independent of Closure 1

Zippering from Closure 2 is not impaired in embryos which lack Closure 1 due to deletion of the core planar cell polarity component *Vangl2* (Fig. 3A-C). These *Vangl2^−/−^* embryos invariably develop craniorachischisis (Fig. 3A-B’’). Nonetheless, their Closure 2 zippering point assembles actomyosin cables (Fig. 3A, A’,B, B’) and in the absence of rostral zippering from Closure 1, zippering can progress from Closure 2 to a more caudal level, at least 100 μm closer to the otic pits than their wild-type counterparts (Fig. 3C). This demonstrates functional redundancy as the rostral closure-initiation point partially compensates to cover a greater proportion of the future mid/hindbrain.

**Figure 3:**
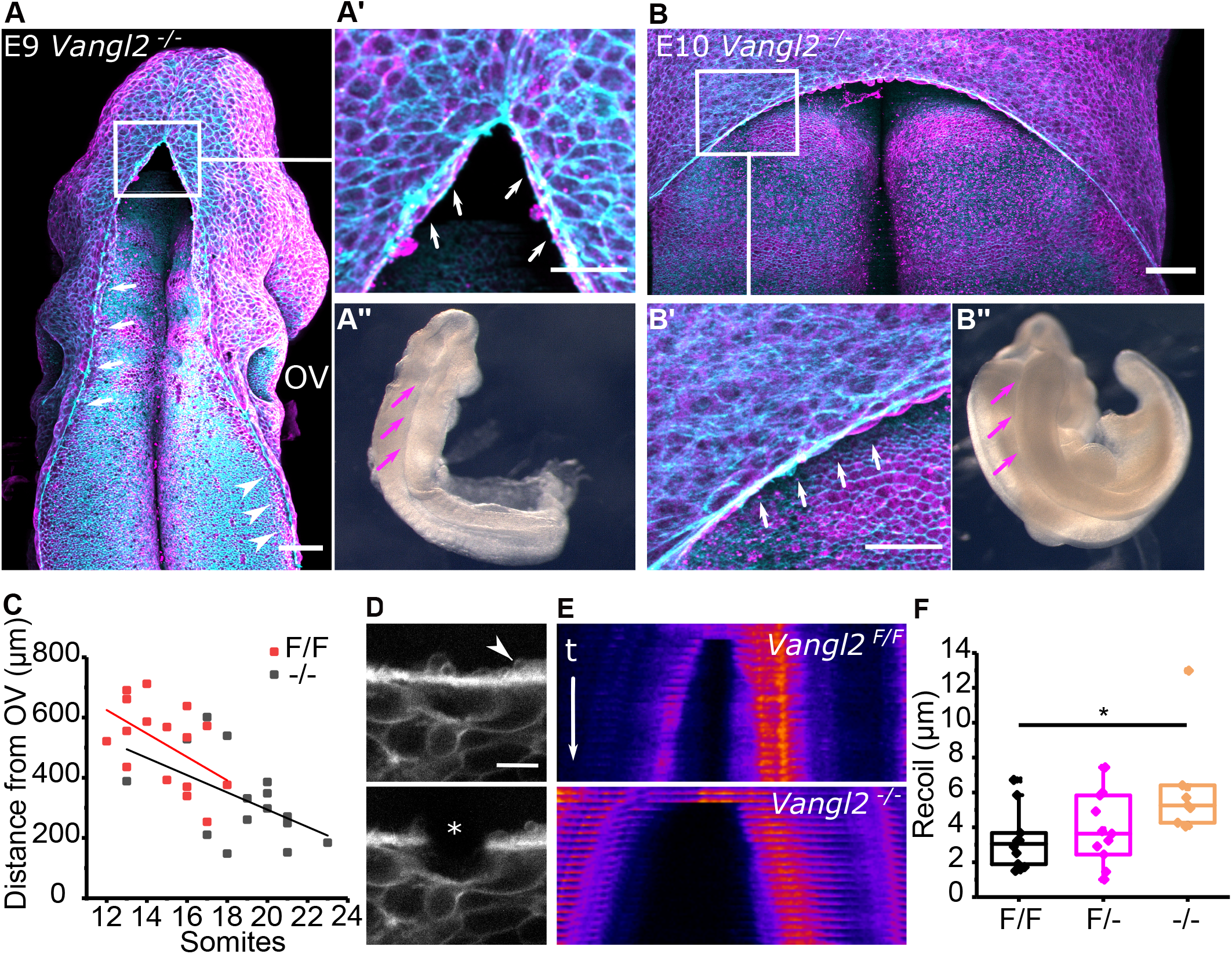
Progression of HNP closure from Closure 2 is independent of Closure 1. **A-A’’:** Representative whole-mount staining and corresponding bright-field image of an early *Vangl2^−/−^* embryo (16 somites). Annotated are the actomyosin cables and associated ruffles extending along the open hindrain (white arrows in **A**, **A’**) and spinal region (white arrowheads in **A**). Magenta arrows indicate the open neural folds in brightfield (**A’’**). Scale bars: 100 μm. **B-B’’:** Representative whole-mount staining and corresponding bright-field image of a late *Vangl2^−/−^* embryo (30 somites). Annotated are the actomyosin cables and associated ruffles extending along the open hindrain (white arrows in **B’**). Magenta arrows indicate the open neural folds in bright-field (**B’’**). Scale bars: 100 μm. **C.** Quantification of the distance between the Clo2 zipper and the mid-OV level in *Vangl2^−/−^* and *Vangl2^F/F^* (control) embryos at the indicated somite stages. The OV level was defined as in Fig 2E. **D.** Representative laser ablation of cable-associated cell borders near the Clo 2 zipper. The asterisk shows the ablated border. The white arrowhead points at membrane ruffles, which co-localise with the cable (see Figure 1D). Scale bar: 10 μm. **E.** Representative kymographs of cable ablations in *Vangl2^F/F^* and *Vangl2^−/−^* embryos. t indicates timeframes post ablation (< 1 s/timeframe) and is the same for both kymographs. **F.** Recoil quantification after cable ablations in *Vangl2^F/F^* (n = 10), *Vangl2^F/−^* (n = 11) and *Vangl2^−/−^* (n = 7) at 12-17 somite stages. P< 0.05, repeated-measures ANOVA with Bonferroni post hoc correction.

Without Closure 1, the actomyosin cables cannot form an encircling purse-string, but instead are present as long cables extending from the Closure 2 zippering point down as far as the unfused spinal neural folds (Fig. 3A, B). Persistent myosin enrichment suggests they are contractile despite this dramatic change in morphology. Confirming this, actomyosin cable tension inferred from recoil after laser ablation is greater in *Vangl2^−/−^* embryos than wild-type littermates (Fig. 3D-F). Cable recoil was assessed in the comparable region adjacent to the Closure 2 zipper at equivalent somite stages. Thus, this transgenic model of extreme HNP asymmetry demonstrates that zippering from rostral and caudal closure points is largely independently of each other.

### Closure rate asymmetry does not arise from local differences in mechanical tension

In wild-type mice, the actomyosin purse-string appears to link the two closure points and differential contractility might explain asymmetric closure rates. Contrary to this, we found that tension withstood by surface ectoderm cell borders engaged in the purse-string cables is comparable between the rostral and caudal extremes of the HNP (Fig. 4A-D). Surface ectoderm cell borders overlaying the closed NT were ablated as comparators, demonstrating approximately five-fold lower recoil compared with cable cell borders after laser ablation (Fig. 4A-D).

**Figure 4.**
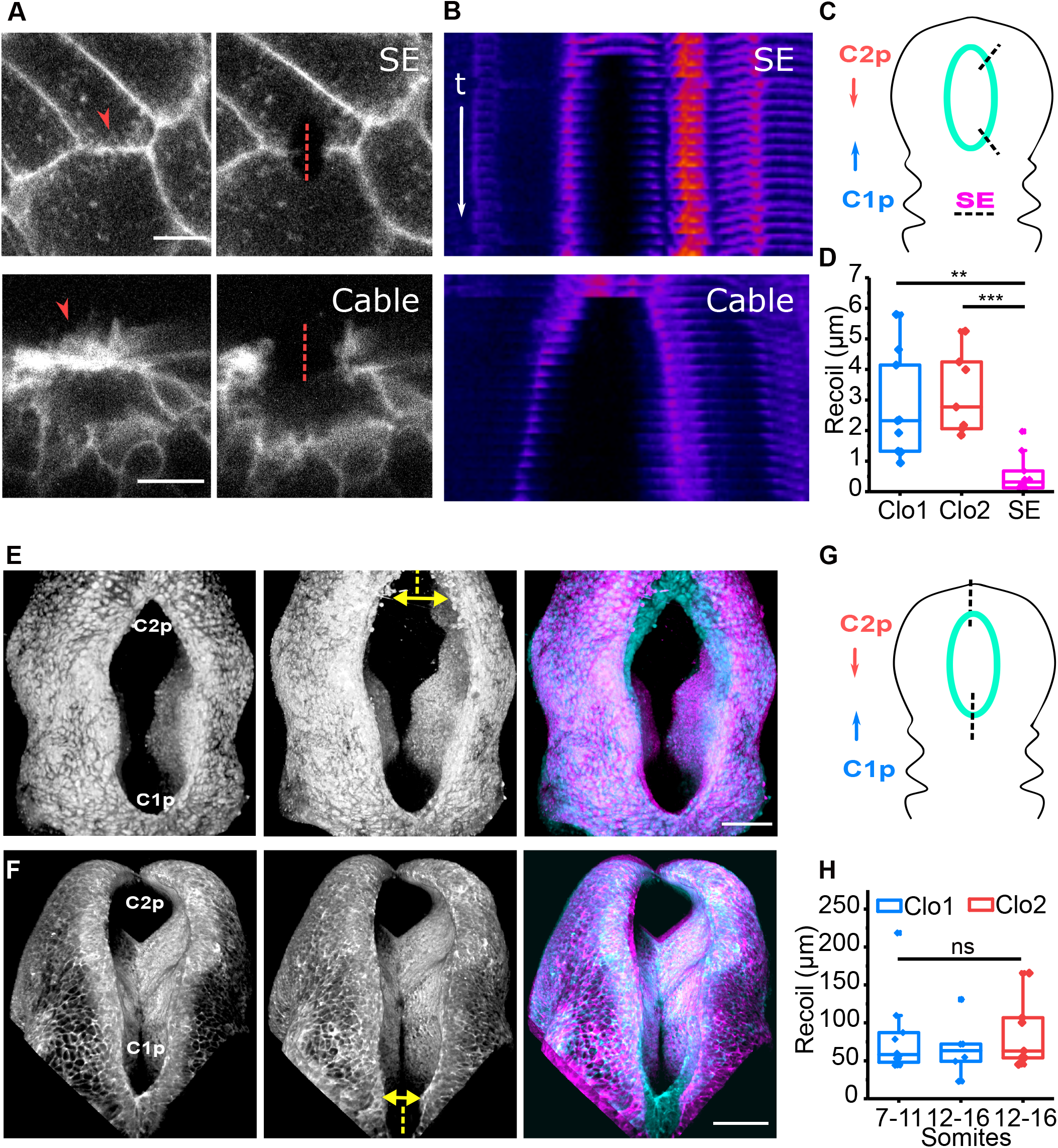
Biomechanical tension is comparable between the rostral and caudal ends of the HNP. **A.** Representative laser ablations of SE border (top) and cable near the Clo2 zipper (bottom). The arrowhead points at membrane ruffles and the dashed line shows the site of ablation. Scale bar: 10 μm. **B.** Kymographs corresponding to the ablations in **A** shown in Fire LUT. t indicates timeframes post ablation (<1 s/timeframe) and is the same for both kymographs. Following bright points (cell features) over time shows faster separation from the ablation site after cable than non-cable surface ectoderm border ablation. **C.** Schematic representation of positions where surface ectoderm (SE) cell border and cable ablations were performed. The HNP rim is indicated in cyan. C1p: closure 1 progression, C2p: closure 2 progression. **D.** Recoil quantification after cable ablations proximal to Clo 1 zipper (n = 9), Clo 2 zipper (n = 7) and ablations of SE borders in the closed region (n = 10). **P< 0.01, ***P<0.001 repeated-measures ANOVA with Bonferroni post hoc correction. **E-F.** 3D reconstruction of representative zippering point (tissue level) laser ablations. Ablations of Clo 2 and Clo 1 zippers are shown in E and F respectively. The dashed line shows the site of ablation and the arrow indicates lateral recoil of the neural folds. Right panel shows the overlay of pre- and post-ablation images in cyan and magenta respectively. Scale bar: 100 μm. **G.** Schematic representation of zippering point (tissue-level) ablations. The HNP rim is indicated in cyan. C1p: closure 1 progression, C2p: closure 2 progression. **H.** Recoil quantification after ablations of the Clo1 zipper at 7-11 somites (n = 10) and 12-16 somites (n = 6) and the Clo2 zipper (n = 7). ns: non-significant, repeated-measures ANOVA with Bonferroni post hoc correction.

In addition to cell border ablations, tissue-level ablations were performed in independent embryos to infer mechanical tension opposing HNP closure as we previously reported in the spinal NT (15–17, 35). Tissue-level laser ablations of either the Closure 1 or Closure 2 zippering points both produced lateral recoil of the neural folds, widening the HNP (Fig. 4E-H). This demonstrates that the recently-fused NT is load-bearing (Fig. 4E-F). HNP widening after tissue ablation also demonstrates that the neural folds are not maintained in apposition by compression from surrounding tissues. We next tested whether tissue-level tension changes as Closure 2 forms, converting the open cranial neural folds into a HNP. The magnitude of lateral recoil at Closure 1 is not significantly different between early developmental stages before Closure 2 forms versus later stages when both zippering points are present (Fig. 4H). Recoil is also comparable when either the Closure 1 or Closure 2 zippering point is ablated in different embryos (Fig. 4H), demonstrating equivalent tissue-level tension at the two extremes.

Thus, neither differential pro-closure contractility (cable tension) nor anti-closure forces (tissue-level lateral recoil) are sufficient to explain asymmetric HNP closure dynamics.

### Combination of cell crawling and purse-string contraction describes HNP closure dynamics

To better understand the mechanistic origin of HNP closure dynamics, and specifically the differential closure rates between Clo1 and Clo2, we developed a vertex-based mechanical model of surface ectoderm cells. Vertex models have been used to describe wound-healing responses of embryonic epithelia, in which physical properties of the tissue substantially influence closure dynamics (36–38). In two-dimensional vertex models, as implemented here, a network of edges represents cell-cell junctions, and the polygons represent the apical surfaces of the cells (Fig. 5A) (39, 40). Each cell has a mechanical energy composed of area elasticity, cytoskeletal contractility, and interfacial tension at the cell-cell junctions. Cell edges lining the HNP gap have an increased tension due to the assembly of the contractile actomyosin purse-string, which generates a driving force for gap closure. Potential models were iteratively tested for their ability to replicate the observed tissue-level HNP closure dynamics, namely an asymmetric closure rate while maintaining an elongated aspect ratio over long timescales.

**Figure 5:**
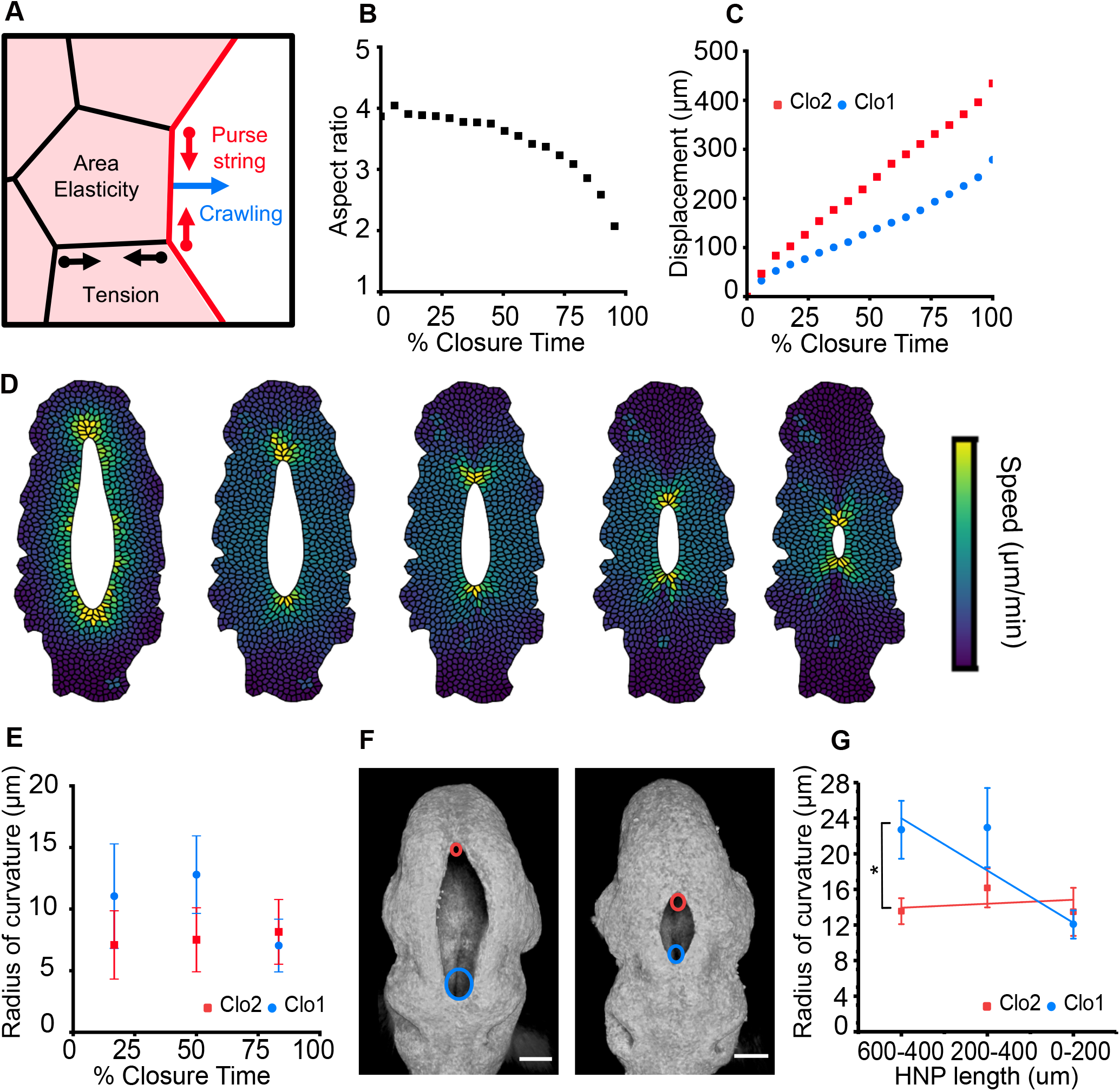
Cell-based modelling reveals that asymmetric geometry regulates closure rate asymmetry. **A.** Schematic of the vertex model for HNP gap closure. Each cell has a preferred area, contractility, and interfacial tension. Cell edges on the gap, highlighted in red, have an increased tension due to the assembly of actomyosin purse-string. Cells around the gap also actively crawl towards it. **B.** Gap aspect ratio against percentage closure time, as outputs of the model combining purse-string contraction and cell crawling. **C.** Displacement of the Closure 1 and 2 zippers against percentage closure time, as outputs of the model. **D.** Time course of simulated gap closure, at 10%, 30%, 50%, 70%, and 90% of closure time, from left to right. Cell colour indicates cell speed. **E.** Simulation mean radius of curvature at the Closure 1 and 2 zippers. Data are binned into the nearest tertile of closure. Error bars represent standard deviation within bins (n = 56). **F.** 3D reconstructions of representative early (left) and late (right) HNPs. A circle was fitted at each zippering point to calculate the radius of curvature. Cyan and red circles annotate closure 1 and 2 zippering points respectively. **G.** Experimental mean radius of curvature at the closure 1 and 2 zippers plotted against HNP length. Error bars represent standard error (n = 14-17 for each bin). P<0.05, paired t test.

Purse-string contractility is nominally sufficient to achieve gap closure, but produces a progressively more circular gap because the zippering points move rapidly towards each other with very little lateral motion (Fig. S1A-D). To maintain an elongated gap shape we included active forces for cell crawling, assumed to occur throughout the bulk of the cell via adhesions onto newly assembled extracellular matrix. We assume that there is always matrix for the cell to crawl on behind their leading edge, and so do not explicitly model the assembly of matrix in the gap. Cell crawling alone is also nominally sufficient to close the gap, but the gap width and length close at the same speed, resulting in increasing gap aspect ratio over time before both sides meet laterally (Fig. S2A-D). Simulating HNP closure from an empirically-determined initial geometry with a combination of purse-string and crawling mechanisms maintains gap aspect ratio over long timeframes (Fig. 5B), closely recapitulating the pattern seen *in vivo* (Fig. 2D).

Varying the speed of cell crawling relative to the default model demonstrates that increasing crawl speeds leads to the maintenance of gap aspect ratio for longer times (Fig. S3A) and increases overall closure rates (Fig. S3B), suggesting that cell crawling is essential for rapid closure of elliptical gaps. Taken together, these simulations suggest that both purse-string contraction and cell crawling contribute to the rate of HNP closure and their combination is necessary to maintain gap aspect ratio over time. We find that asymmetry between rostral and caudal closure rates is greater when purse-string contraction is the only driving force (Fig. S1B), is present when cell crawling is the only closing force (Fig. S2B), and continues to be observed when both purse-string contraction and cell crawling are implemented (Fig. 5C). This asymmetry in closure rate arises despite equal cable tension or crawling forces at Clo1 and Clo2.

Recent work has suggested that gap geometry plays an important role in regulating the dynamics of wound closure, such that the rate of purse-string driven closure is proportional to the gap curvature (36, 41, 42). Indeed, the HNP geometry substantially differs between the rostral and caudal regions (Fig. 1B, D, Fig 2A), motivating us to investigate the relationship between closure rate asymmetry and HNP geometry.

### Tissue geometry produces closure rate asymmetry

Simulated closure of an arithmetically elliptical gap, rather than empirically determined HNP geometry, produces equivalent rates of closure from the two extremes (Fig. S4A, B). For a rate asymmetry to exist, there must be some difference between the two closure points in either the driving forces arising from purse-string and cell crawling, or the resistive forces arising from the surrounding tissue. While the tension was measured to be equivalent around the gap (Fig 4), gap geometry differs. We therefore interrogated components of local tissue properties in addition to purse-string tension and cell crawling in the model. At early stages there are fewer cells in the plane of the gap in the recently closed region rostral to Closure 2 than there are in the closed region caudal to Closure 1 (see Fig. 1B). Closure of an elliptical gap on biologically realistic boundary conditions progresses slightly faster from the end with fewer cells (Fig. S4C, D). Exaggerating differences in boundary conditions increases closure rate asymmetry (Fig. S4E, F). Thus, an ellipse can close asymmetrically when fewer cells need to be re-arranged at one apex than the other (Fig. S4G), but this effect is insufficient to explain the observed closure rate asymmetry under biologically relevant conditions. The number of cells surrounding the gap minimally affects the maintenance of gap aspect ratio during ellipse closure (Fig. S4H).

An additional geometric feature incorporated in the simulation is that, in early embryos with long HNPs, the Clo1 zippering point has a greater radius of curvature than that at Clo2 (Fig. 5E-G). In both the simulation (Fig. 5E) and *in vivo* (Fig. 5F, G), Clo2 maintains a low radius of curvature (more acute angle), while Clo1’s radius of curvature decreases over developmental time. Since the purse-string acts as a cable under tension, the resulting force will be inversely proportional to the radius of curvature, resulting in a larger net force and faster closure at Closure 2 (Fig. S1B). Consequently, simulated purse-string driven closure displays gap length-dependant dynamics: it slows down as the HNP becomes rounder, before speeding up again when the gap is very small (Fig. S5).

The rate of shortening is more constant throughout the closure period when crawling is implemented alone or in addition to purse-string contraction (Fig. S5). A constant shortening rate is more consistent with the overtly linear relationship between HNP length and somite stage observed *in vivo* (Fig. 2B). Closure rate remains faster from Closure 2 when cell crawling is the only driving force, despite cells crawling at the same speed (Fig. S2B). This is because the gap is narrower near the Closure 2 zipper, so the edges meet sooner. Incorporating this experimentally determined gap geometry in the final model reproduces the asymmetry in closure rates observed *in vivo* (Fig. 5C-D).

### Surface ectoderm cells display HNP gap-directed crawling *in vivo*

Having developed an *in silico* model which meaningfully recapitulates tissue-level dynamics, we used it to predict the underlying cell-level dynamics around the HNP rim. In particular, we investigated the dynamics of three rows of cells around the HNP, with Row 1 being the most proximal cells which form the actomyosin purse-strings (Inset in Fig. 6A). The three rows describe concentric rings with progressively more cells in each row. During simulated HNP closure, the number of cells in each row decreases gradually, with row occupancy decreasing at the same rate in each row (Fig. 6A). Cells nearest to the gap move with greater speed during closure (Fig. 6B) but have slightly lower directionality (defined as Euclidian distance divided by total distance travelled) compared to the surrounding rows (Fig. 6C-D).

**Figure 6:**
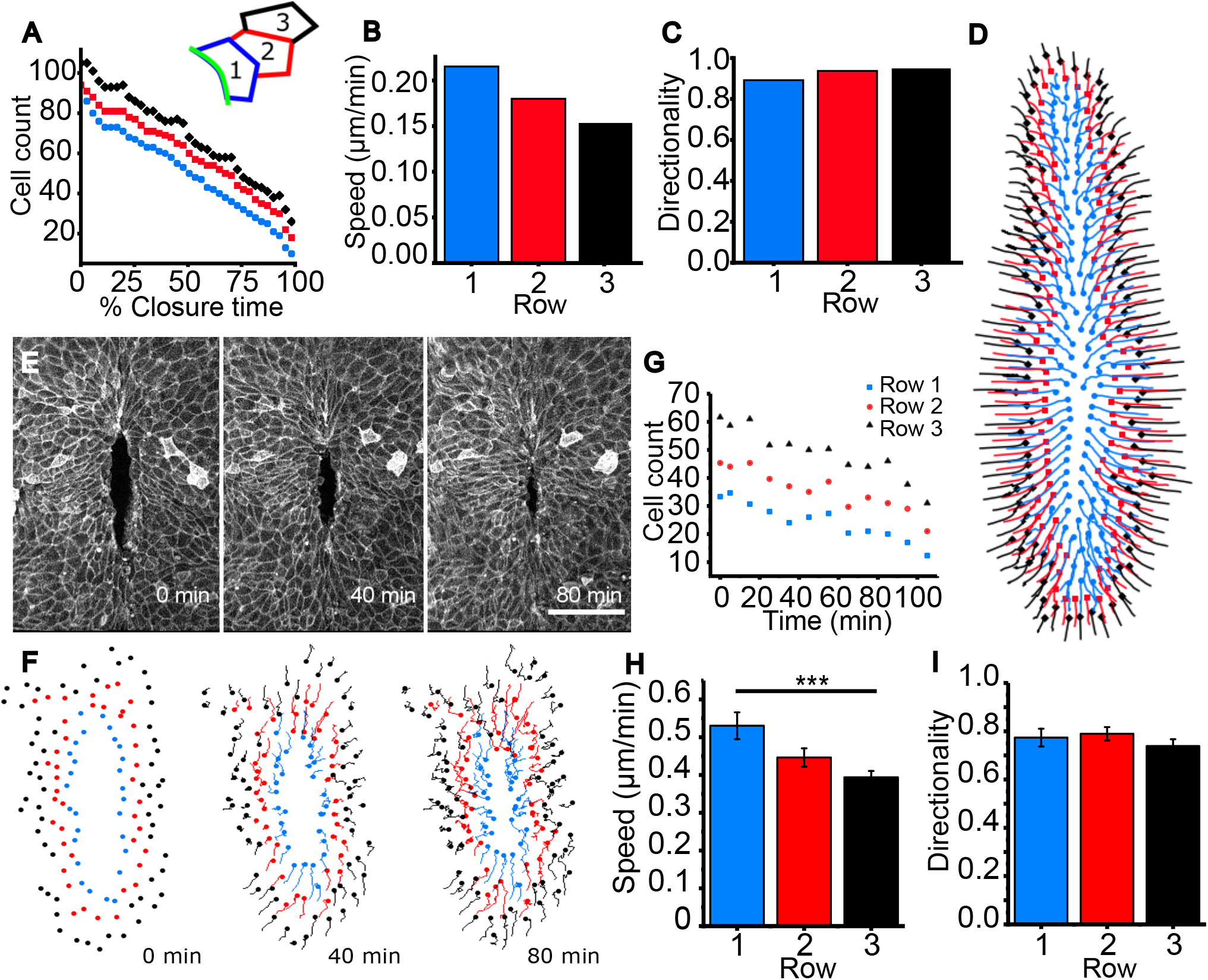
Directional migration of surface ectoderm cells towards the HNP gap. Data in panels A-D are from simulated gap closure achieved by combined purse-string constriction and cell crawling, whereas E-I are from live-imaged embryos. **A.** Cell count in the first 3 rows around the gap against closure time. The inset illustrates the three cell rows analysed. Row 1 cells (blue) assemble the cables (green line) at the HNP rim. Row 2 cells (red) contact Row 1 without being engaged in the cable, Row 3 (black) contact Row 2 cells. **B-C**. Mean speed (B) and mean directionality (**C**) per row. Mean cell speed decreases in rows further from the simulated gap, but directionality remains comparable between rows. **D.** Cell center trajectories over the course of closure. Colour indicates initial cell row. **E.** Snapshots of a live imaged mTmG mouse embryo at the time points indicated showing progression of HNP closure at sub-cellular resolution. Scale bar: 100 μm. **F.** Tracks of individual cells at the first three rows around the HNP shown in E, illustrating directional cell crawling during live imaging. **G.** Number of cells per row over time of the embryo live-imaged in E showing equivalent rates of row occupancy. **H.** Mean cell speed per row for the first 25 min, showing Row 1 cells underwent faster displacement than Rows 2 or 3 in this embryo. P<0.001, One-way ANOVA with Bonferroni post-hoc correction. **I.** Mean cell directionality per row for the first 25 min, showing equivalence between rows.

We find similar trends in models with only cell crawling, or only purse-string contraction. The velocity is highest, and directionality low, in the cells closest to the gap (Purse-string only Fig. S1E, F; Crawling only Fig. S2E, F). However, the directionality of Row 1 cell migration is much lower (< 0.7 Euclidian/accumulated distance) when purse-string mediated closure is implemented without cell crawling (Fig. S1F) because cells display large curving trajectories (Fig. S1C).

Surface ectoderm crawling, to our knowledge, has not previously been reported to drive HNP closure. We therefore developed sub-cellular live-imaging capability in order to visualize surface ectoderm rearrangement around the closing HNP (Videos S2, 3). Manual tracking of surface ectoderm cells in the first three rows confirmed they crawl towards the HNP gap (Fig. 6E-F, Video S4). As predicted by our vertex model, the rates at which cells leave each row is equivalent between rows (Fig. 6G, representative of 4/4 independently live-imaged embryos). Cell migration parameters were analysed over the first 25 minutes of live imaging to reduce re-analysis of cells which switch rows. Over this period, cells in Row 1 had a significantly higher speed than Row 3 in 2/4 embryos (representative in Fig. 6H) and an equivalent speed in the remaining 2/4 embryos. Row 1 cells had equivalent (3/4 embryos, Fig. 6I) or significantly higher (1/4 embryo) directionality than row 3 cells. Directionality of row 1 cells was > 0.7 Euclidian/accumulated distance in all four independent embryos, arguing against cell displacement due to purse-string constriction which produces lower directionality in our simulations (Fig. S1F). Taken together, live imaging confirms that highly directional surface ectoderm cell crawling towards the gap contributes to HNP closure.

## Discussion

The interdisciplinary studies described here establish a conceptual biophysical framework through which disruption, or enhancement, of HNP closure can be assessed. Tissue geometry, directional surface ectoderm crawling and actomyosin purse-string contractility emerge as necessary parameters sufficient to describe the simulated dynamics of HNP closure. Each of these three parameters will be responsive to a large number of genes and signalling cascades, whose enhancement may promote timely closure and disruption may impede closure. Simulation demonstrates both purse-string contractility and cell crawling accelerate closure at Closure 2 due to its lower radius of curvature observed *in vivo*. This means a greater proportion of the mouse HNP roof plate is produced by zippering from Closure 2 because tissue geometry constrains progression from Closure 1. Moreover, our model is able to capture the directional dynamics of the surface ectoderm cells surrounding the gap visualized by live imaging.

Model systems used to study gap closure include *in vitro* wounds (37, 43–45), *Drosophila* dorsal closure (46, 47), embryonic wound healing (48, 49), nematode ventral closure (50), and *Xenopus* blastopore closure (51, 52). Collectively, these systems have established cellular and biomechanical mechanisms commonly employed to close gaps. Recurring mechanisms include the formation of contractile actomyosin cables at tissue interfaces, cell migration and partner exchange across the gap midline (47, 53, 54). However, differences in gap geometry and cell types involved preclude direct extrapolation of mechanisms identified in other systems to closure of the mammalian NT.

A previously-described mechanism of HNP closure is the establishment of cell contacts across the midline by surface ectoderm cell protrusions (23, 24). We now find these protrusions appear to emanate from the actomyosin purse-strings surrounding the HNP rim. Actomyosin purse-strings are robust morphogenetic tools which couple cell-level control of morphogenesis to tissue-level deformation (17, 55, 56). F-actin polymerisation and turnover mechanisms by which these structures are assembled and maintained in the HNP remain to be determined: global deletion of F-actin regulators including formin homology 2 domain-containing 3 (57), shroom-3 (27, 58) and cofilin-1 (59, 60) each preclude HNP formation. The requirement for actomyosin purse-strings in gap closure has recently been questioned in *Drosophila* because dorsal closure completes successfully in embryos lacking high-tension purse-strings (61, 62). However, there are substantial differences between these systems: the early HNP is approximately double in length and takes three times as long to close as the *Drosophila* dorsal gap. Dorsal closure is slower in fly embryos which lack purse-strings (61, 62), suggesting that one of the roles of these structures in the HNP could be to ensure closure completes before the previously-suggested (14) developmental deadline.

The importance of seemingly subtle tissue geometric properties in determining HNP closure rate have not, to our knowledge, previously been appreciated. The inward closure force generated by the purse-string is greater in regions of higher rim curvature (42), increasing cell speeds at the Closure 2 zippering point. In addition, crawling cells have less distance to cross at the Closure 2 compared to Closure 1 zipper. Why the Closure 1 zippering point is wider than that at Closure 2 in the early HNP is unclear. Potential explanations include very different mechanisms underlying the initial formation of these closure points and whole-embryo deformation during axial rotation after Closure 1 forms (63). Intriguingly, the initial position at which Closure 2 forms varies between genetically wild-type mouse strains (64), suggesting a degree of functional redundancy. Morphogenetic redundancy, or compensation, is demonstrated in this study by the ability of the Closure 2 zippering point to proceed further caudally in the absence of Closure 1.

Embryos with craniorachischisis studied here also demonstrate that the assembly of high-tension actomyosin supracellular enrichments does not require Closure 1 or expression of *Vangl2*. This is consistent with a previous report that *Vangl2* deletion does not impair wound healing in the foetal epidermis (65). Increased actomyosin purse-string tension in *Vangl2^−/−^* embryos compared with littermate controls could either indicate *Vangl2* normally suppresses contractility or, more likely, may be secondary to tissue-level structural differences. It is well-established that mechanical tensions triggers mechano-chemical feedback mechanisms which increase non-muscle myosin recruitment (66) and adherens junction stability (67).

Establishment of E-cadherin adherens junctions between surface ectoderm cells on opposite sides of the embryonic midline allows advancement of the zippering points. The zippering dynamics observed during live-imaged HNP closure in this work is different from what has been described in other closure processes. Individual surface ectoderm cells at the zipper appear to follow this point, often migrating at the leading edge for over one cell length. In contrast, live-imaging of *Ciona* NT closure shows shrinkage of cell junctions ahead of the zipper and direct matching of cells across the embryonic midline (68). Although a “buttoning” method had previously been suggested to close the HNP based on lower-resolution live-imaging (69), no evidence of “buttoning” protrusions are observed in freshly-dissected embryos fixed directly after removal from the uterus, nor are they visible in the high-resolution live imaging provided here.

Previous live-imaging of mouse HNP closure also demonstrated displacement of cell nuclei at the leading edge of the gap (70). The first row of surface ectoderm cells extends protrusions toward the neuropore, similarly to closure of large gaps in wound healing (71, 72). Stochastic lamellipodia-driven migration of a small number of leader cells results in ‘rough’ edges of the closing gap (54). In contrast, the closing HNP gap has ‘smooth’ edges, presumably due to co-ordinated actomyosin cable constriction in row 1 cells. It remains to be established how directionality is inferred in row 1 cells and whether cells at rows 2 and 3 also actively crawl or are passively dragged.

In summary, the biophysical framework presented here begins deconstructing cellular mechanisms of HNP closure from morphometric measurements. Our findings extend the generalisability of core pro-closure modules beyond the size and time scales commonly studied in simpler organisms. Their concurrence encourages generalisation to other closure events of both scientific and clinical importance.

## Materials and Methods

### Animal Procedures

Studies were performed under the regulation of the UK Animals (Scientific Procedures) Act 1986 and the Medical Research Council’s Responsibility in the Use of Animals for Medical Research (1993). C57BL/6 mice were bred in-house and used as plug stock from 8 weeks of age. Mice were mated overnight and the next morning a plug was found and considered E0.5. In some cases, mice were mated for a few hours during the day and the following midnight was considered E0.5. Pregnant females were sacrificed at E8.5 (~12 somites) or E9 (~17 somites). *Vangl2^Fl/−^* mice were as previously described (Ramsbottom et al., 2014) and were always phenotypically normal. To obtain *Vangl2^−/−^* embryos, *Vangl2^Fl/−^* stud males were crossed with *Vangl2^Fl/−^* females. *Vangl2^Fl/Fl^* embryos were used as littermate controls. mTmG mice were as previously described (Muzumdar et al., 2007) and tdTom fluorescence from homozygous mTmG embryos was used for live-imaging.

### Immunofluorescence, image acquisition and analysis

Embryos were dissected out of their extraembryonic membranes, rinsed in ice-cold PBS and fixed in 4% PFA overnight (4°C). Whole-mount immunostaining and imaging were as previously described (Galea et al., 2017). Primary antibodies were used in 1:50-1:100 dilution and were as follows: rabbit E-cadherin (3195, Cell Signalling Technology), mouse N-cadherin (14215S, Cell Signalling Technology), goat fibronectin (SC-6952 (C-20), Santa Cruz Biotechnology) and rabbit MHC-IIB (909901, BioLegend). For N-cadherin staining, antigen retrieval was first performed for 1 h at 100°C, using 10 mM sodium citrate with 0.05% Tween 20, pH 6.0. Secondary antibodies were used in 1:200 dilution and were Alexa Fluor-conjugated (Thermo Fisher Scientific). Alexa Fluor-568-conjugated Phalloidin was from Thermo Fisher Scientific (A121380). Images were captured on a Zeiss Examiner LSM 880 confocal using 10 x/NA 0.5 or 20 x/NA 1.0 Plan Apochromat dipping objectives. Whole HNP images were captured with x/y pixel sizes of 0.42-0.59 μm and z-step of 0.8-2.44 μm (speed, 8; bidirectional imaging, 1024×1024 pixels). Images were processed with Zen 2.3 software and visualised as maximum projections in Fiji (Shindelin et al., 2012). To visualise the surface ectoderm, the z-stacks were first surface subtracted as previously described (Galea et al., 2018) to only show the apical 2-3 μm of tissue (macro available at https://www.ucl.ac.uk/child-health/core-scientificfacilities-centres/confocal-microsopy/publications).

For morphometric analysis, HNP length and width were calculated by annotating the HNP rim and then measuring the major and minor axis using the fit ellipse function in Fiji. To quantify the distance of each zipper from the otic vesicles, reflection images were captured using the 10 x/NA 0.5 dipping objective (633 nm wavelength, x/y pixel size 2.44, z step 3.33 μm). The z stacks were 3D rotated and visualised as maximum projections. For 3D visualisation of reflection images (Fig 2A, E), z-stacks were despeckled in Fiji, filtered with a Kuwahara filter (sampling window width of 5) and opened with the 3D viewer plugin.

### Live imaging

Live imaging was performed as previously described (Galea et al., 2017, Mole at al., 2020). Embryos were dissected with an intact yolk sac and transferred into 50% rat serum in DMEM. They were then held in place with microsurgical needles (TG140-6 and BV75-3, Ethicon) and a small window was made in the yolk sac and amnion, exposing the HNP. Heart beat was steady throughout each experiment. Images were captured on Zeiss Examiner LSM 880 confocal (37°C, 5% CO_2_), using a 20x/NA 1.0 Apochromat dipping objective. X/Y pixels were 0.27-0.83 μm and z step was 1 μm. The time step was 5-10 min. Five embryos from independent litters were live-imaged for a minimum of 1 hour each.

Live imaging datasets were 3D registered in Fiji using the Correct 3D Drift plugin (Parslow et al., 2014). They were then deconvolved using the Richardson-Lucy algorithm (5 iterations) in DeconvolutionLab2 (Sage et al., 2017). All sequences were surface subtracted (macro above) in order to enable visualisation of surface ectoderm cell borders. Cell migration analysis was done in Fiji using the manual tracking plugin along with the ‘chemotaxis and Migration tool’ plugin (ibidi). Particle Image Velocimetry (PIV) analysis was performed in Fiji using the in-built Iterative PIV (Cross-correlation) plugin (32 pixel final interrogation window size, normalise median test noise = 1 and threshold = 5). Images were Gaussian-filtered (radius = 2 pixels) before applying PIV.

### Laser ablations

After removal of the extraembryonic membranes, embryos were stained with 1:500 CellMask Deep Red plasma membrane (C10046 Invitrogen) in DMEM at 37°C for 5 min. They were then positioned on agarose plates using microsurgical needles and moved to the microscope stage (heated at 37°C). Tissue-level (Galea et. al., 2019) and cable (Butler et al., 2019) laser ablations were performed as previously described using a Mai Tai laser (SpectraPhysics Mai Tai eHP DeepSee multiphoton laser).

For cable ablations, a 0.1 μm-wide line was cut using 710 nm wavelength at 100% laser power (0.34 μs pixel dwell time for 10 iterations, 20X/NA 1 Plan Apochromat dipper). One ablation was analysed per embryo. Cable recoil was calculated by measuring the immediate displacement of cell landmarks perpendicular to the ablation.

For tissue-level zippering point ablations, a pre- and post-ablation z-stack was obtained using 10X magnification at 633 nm. Total acquisition time for each stack was ~3 min. The ablations were performed using 800 nm wavelength at 100% laser power (65.94 μs pixel dwell time for 1 iteration, 10X/NA 0.5 Plan Apochromat dipper). The zippering point was ablated using narrow rectangular ROIs, moving sequentially in z to ensure the tissue was ablated.

### Statistical Analysis

All statistical analysis was performed in OriginPro 2017 (Origin Labs). Individual embryos were the unit of measure. Images are representative of embryos from a minimum of three independent litters. Comparison of two groups was by Student’s t-test, paired by embryo where appropriate. Comparison of multiple groups was by one-way ANOVA with post-hoc Bonferroni. Graphs were made in OriginPro 2017 and are shown either as box plots or as mean +/− SEM, when several embryos were averaged per data point. For box plots, the box shows the 25-75^th^ percentiles and the median is indicated by a line. The whiskers show the 95% confidence intervals and the outliers are indicated (not excluded).

In Fig 2 F, G, distances of the Clo1 and 2 zipper were normalized against the longest distances recorded for each zipper. Linear regression slopes were estimated using Pearson’s regression and compared by F-test.

### Computational Model

To model neural tube closure, we use the vertex model for epithelia (39, 40). The apical surface of the tissue is modelled by a network of connected edges, with cells described as the polygons and cell-cell junctions as the edges. The tissue mechanical energy given by:

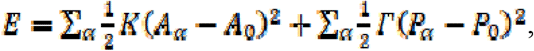

where *α* indicates the sum over all cells. The first term represents cell area elasticity, with elastic modulus *K*, cell area *A_α_* and preferred area *A_0_*. The second term represents a combination cytoskeletal contractility and interfacial adhesion energy, where *Γ* is the contractility, *P_α_* the cell perimeter, and *P_0_* the preferred perimeter. When adhesion dominates over contractility, *P_0_* will be large as cells aim to increase contact length with their neighbours. The mechanical force acting on vertex *i* is given by *F_i_* = −*∂E*/*∂x_i_*, where *x_i_* is the position of the vertex. Assuming that the system is over-damped, the equation of motion is given by:

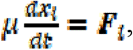

where *μ* is the drag coefficient.

To model the closure forces acting on the border cells at the gap, we implement an increased tension *A_ps_* for edges on the gap representing the purse-string (36, 38). The purse-string tension is chosen such that the total tension of the junction is equal to 5 times the mean tissue tension. The tension within the tissue is given by 2Γ(*P* − *P*_0_), where is the mean cell perimeter, since two cells contribute to a single junction, and at the wound edge by Γ(*P* − *P*_0_) + ⋀_*ps*_. Thus, we calculate ⋀_*ps*_ =9Γ(*P* − *P*_0_). Each cell around the gap may also crawl, with direction 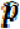 the unit vector perpendicular to their edge on the gap (36). The crawling applies an additional force to all vertices of that cell equal to 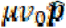, where *ν*_0_ is the cell crawl speed.

### Model implementation

The model is implemented using Surface Evolver (73). We generate an initial tissue configuration using data from experiments. Given a set of cell centers, and boundary points for the gap and border of the tissue, we generate a Voronoi diagram, giving us the cell shapes. The tissue is then relaxed, with the gap edges fixed to maintain its shape, to a mechanical equilibrium before simulating closure. In experiments, the tissue curves around but is free to deform, thus we use free boundary conditions on the external edges. If an edge shrinks below a critical value, *L*_*T*1_, the edge undergoes a rearrangement, or T1 transition, in which a new edge is formed perpendicular to the original junction. The equations are solved numerically by discretizing the equation of motion: *x_i_*(*t*+*Δt*) = *x_i_*(*t*)+*F_i_*/*μ*, where *Δt* is the time step.

### Model parameters

We non-dimensionalise length by 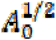 and energy by 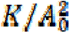, giving us a normalized mechanical energy of

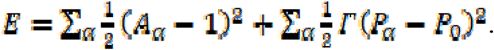

The value of contractility *Γ*, and preferred perimeter *P*_0_, are chosen to minimise the mean square displacement between the initial vertex positions from experiments and final positions after relaxation to equilibrium. Since we are interested in the percentage of closure time, we use a non-dimensional time by rescaling with the drag coefficient *μ*, and choose a small time step for numerical stability.

The purse-string tension chosen so that the total tension on the gap edges is five times greater than the mean edge tension, to match experimental recoil rates after laser ablation. The tension on an edge has contributions from the two cells connected to it, giving a mean tension of 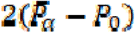, where 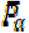 is the mean cell perimeter. Since the gap has contributions from one cell, the purse-string tension satisfies 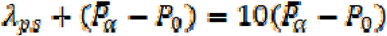. Cell crawl speed is chosen to maintain a constant gap aspect ratio over time.

## Supporting information

Graph source data

## Data availability

The computer code produced in this study is available in the following databases: Vertex Model simulations for HNP Gap Closure: Github (https://github.com/BanerjeeLab/HNP)

## Acknowledgements

This study was funded partly by a Wellcome Trust Postdoctoral Clinical Research Training Fellowship (107474/Z/15/Z) and partly by a Wellcome Clinical Research Career Development Fellowship (211112/Z/18/Z), both to GLG. AM was supported by a Child Health Research CIO award (to NDEG). AJC and NDEG are supported by Great Ormond Street Hospital Children’s Charity. Research infrastructure within the institute is supported by the NIHR Great Ormond Street Hospital Biomedical Research Centre. The views expressed are those of the authors and not necessarily those of the NHS, the NIHR or the Department of Health. MFS is supported by an EPSRC funded PhD studentship at UCL and in part by the National Science Foundation under Grant No. NSF PHY-1748958, and SB acknowledges funding from the Royal Society (URF/R1/ 180187) and HFSP (RGY0073/2018). We thank Dr Dale Moulding for his help with image processing, as well as Dawn Savery and the biological services staff for help with transgenic colonies.

## Author contributions

Conceptualisation: G.L.G., S.B., E.M., M.F.S., N.D.E.G., and A.J.C., Data curation and Formal analysis: E.M., M.F.S., A.M., Investigation: G.L.G., E.M., M.F.S., A.M., Methodology: G.L.G., S.B., E.M., M.F.S., Writing: G.L.G., S.B., E.M., M.F.S. N.D.E.G., and A.J.C, Funding acquisition and supervision: G.L.G. and S.B.

