## Supplementary Figures for "Hindbrain neuropore tissue geometry determines asymmetric cell-mediated closure dynamics"

**Appendix**

**Table of contents (TOC)**

**Appendix Figure S1-** Page 2

**Appendix Figure S2**- Page 3

**Appendix Figure S3-** Page 4

**Appendix Figure S4**- Page 5

**Appendix Figure S5**- Page 6

**Appendix Videos legends**- Page 7

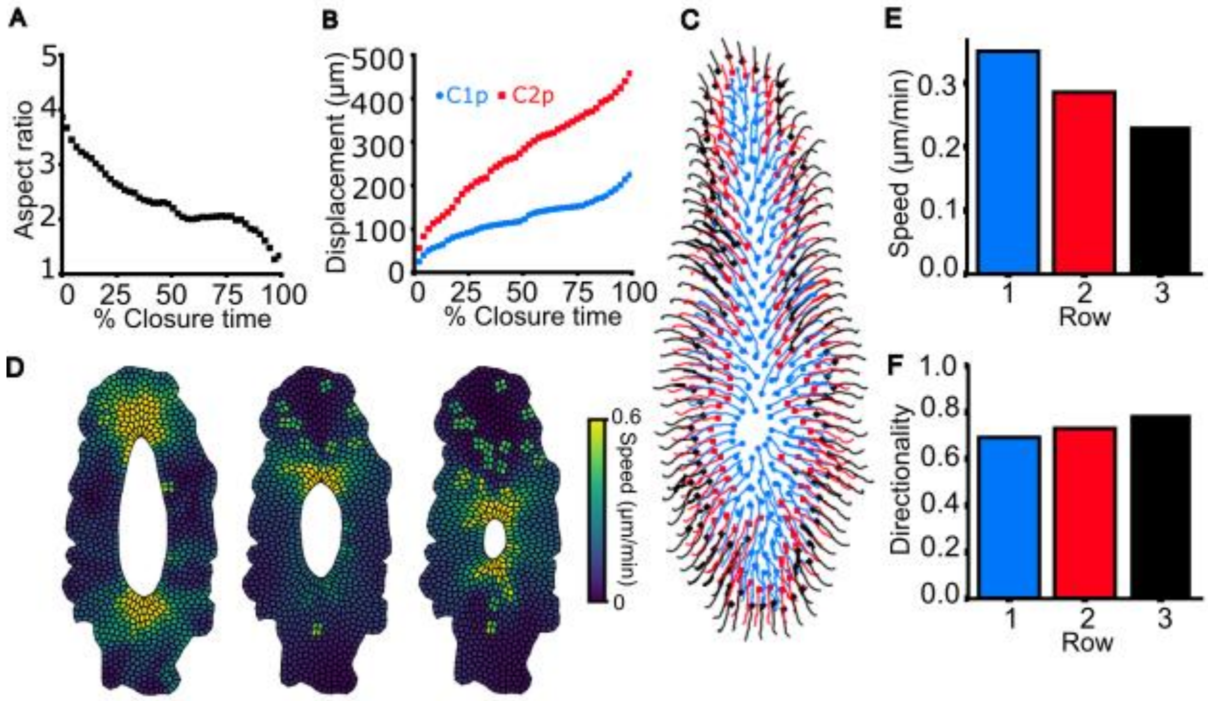


**Appendix Figure S1: Simulations of gap closure with no cell crawling.**

**A-B**. Gap aspect ratio (A), and displacement (B) of Closure 1 and 2, against percentage closure time.

**C**. Cell center trajectories during closure. Colour indicates initial cell row.

**D**. Simulation images over time, showing 10%, 50%, and 90% of closure time, from left to right. The cell colour indicates the cell speed.

**E-F**. Mean velocity (E), and mean directionality (F) for cells in each of the first three rows around the HNP.

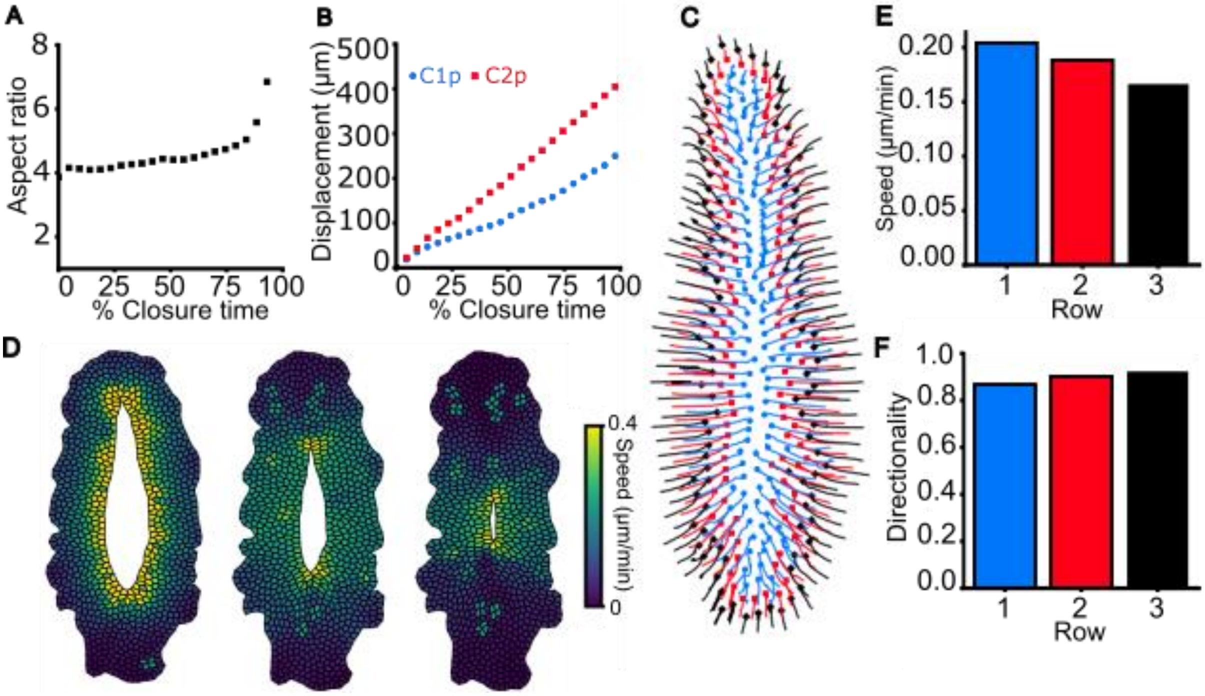


**Appendix Figure S2: Simulation of gap closure with no purse-string.**

**A-B**. Gap aspect ratio (A), and displacement (B) of Closure 1 and 2, against percentage closure time.

**C**. Cell center trajectories during closure. Colour indicates initial cell row.

**D**. Simulation images over time, showing 10%, 50%, and 90% of closure time, from left to right. The cell colour indicates the cell speed.

**E-F**. Mean velocity (E), and mean directionality (F) for cells in each of the first three rows around the HNP.

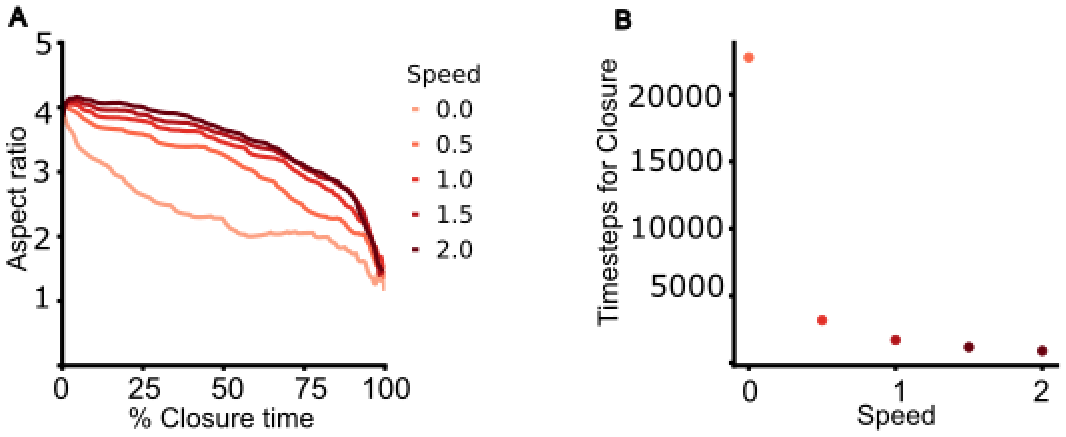


**Appendix Figure S3: Cell crawling maintains an elongated gap aspect ratio and reduces the time taken to complete closure.**

**A-B.** Aspect ratio against percentage closure time (A), and simulation timesteps until closure (B), for different cell crawl speeds, relative to the default case ($v_0$ = 0.01).

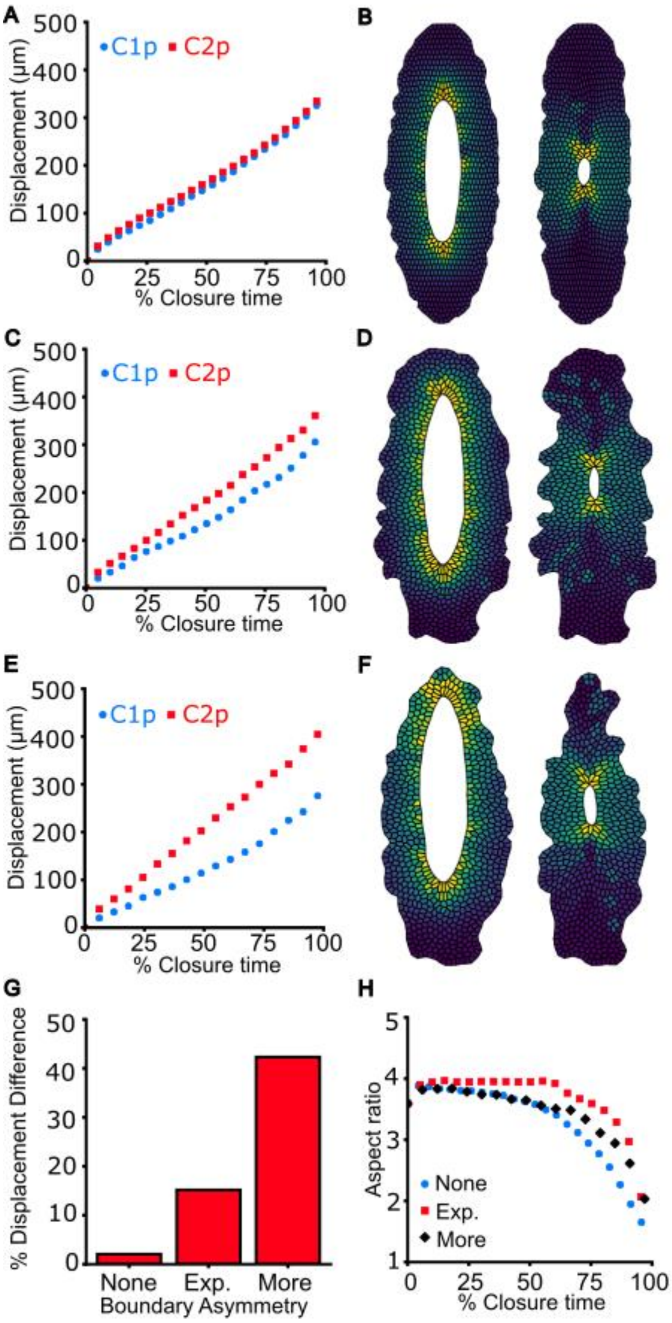


**Appendix Figure S4: Tissue geometry regulates closure rate asymmetry for symmetric gap shapes.**

**A-F**. Displacement of Closure points 1 and 2 plotted against percentage closure time and corresponding simulation snapshots (at 10% and 90% closure time) for symmetric tissue geometry surrounding the gap (A-B), experimental tissue shape (C-D), and a highly asymmetric tissue geometry (E-F).

**G**. Percentage difference between Closure 2 and Closure 1 displacements for the different surrounding tissue geometries.

**H**. Gap aspect ratio against percentage closure time for different geometries of the tissue surrounding the gap.

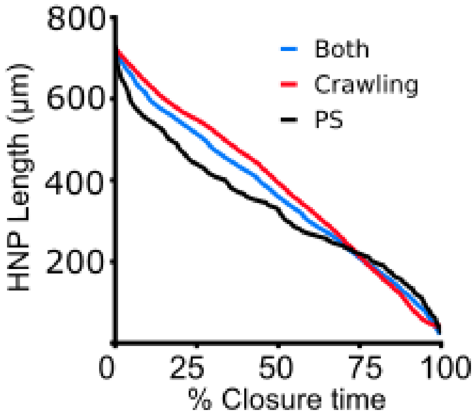


**Appendix Figure S5: Solely purse-string driven closure produces differential closure rates during the closure period.**

HNP length against percentage closure time for simulations with both cell crawling and purse-string, crawling only, and purse-string only.

**Appendix Videos**

**Appendix Video S1: High resolution imaging of surface ectoderm cells on a fibronectin-containing matrix which extends to the HNP rim.** Representative 3D reconstruction of an AiryScan-imaged rostral zippering point showing fibronectin (magenta) extends to the HNP rim, underlying surface ectoderm cells labelled with E-cadherin (green).

**Appendix Video S2: Live imaging of an E9 mTmG homozygous embryo showing the closing HNP**. Clo 2 zipper is at the top and Clo 1 zipper at the bottom of the image. A z-stack was captured every 10 min for a total time of 170 min. Scale bar: 100 µm.

**Appendix Video S3: Live imaging of an E9 mTmG homozygous embryo showing the closing HNP.** Clo2 zipper is at the top and Clo 1 zipper at the bottom of the image. A z-stack was captured every 5 min for a total time of 105 min. Scale bar: 100 µm.

**Appendix Video S4: Manual tracking of the first three rows of cells around the HNP in Video 3.** Cells were tracked for the first 25 min of the movie and move collectively in an inward direction.
