## Supplementary figures and images for "Hindbrain neuropore tissue geometry determines asymmetric cell-mediated closure dynamics"

### Movie S1

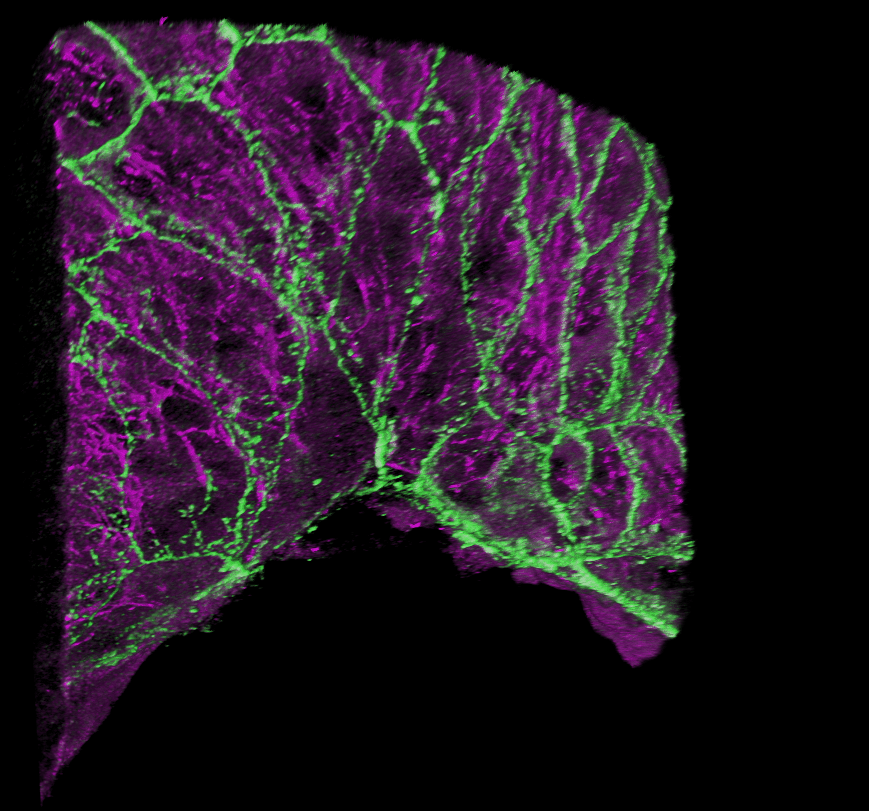

### Movie S2

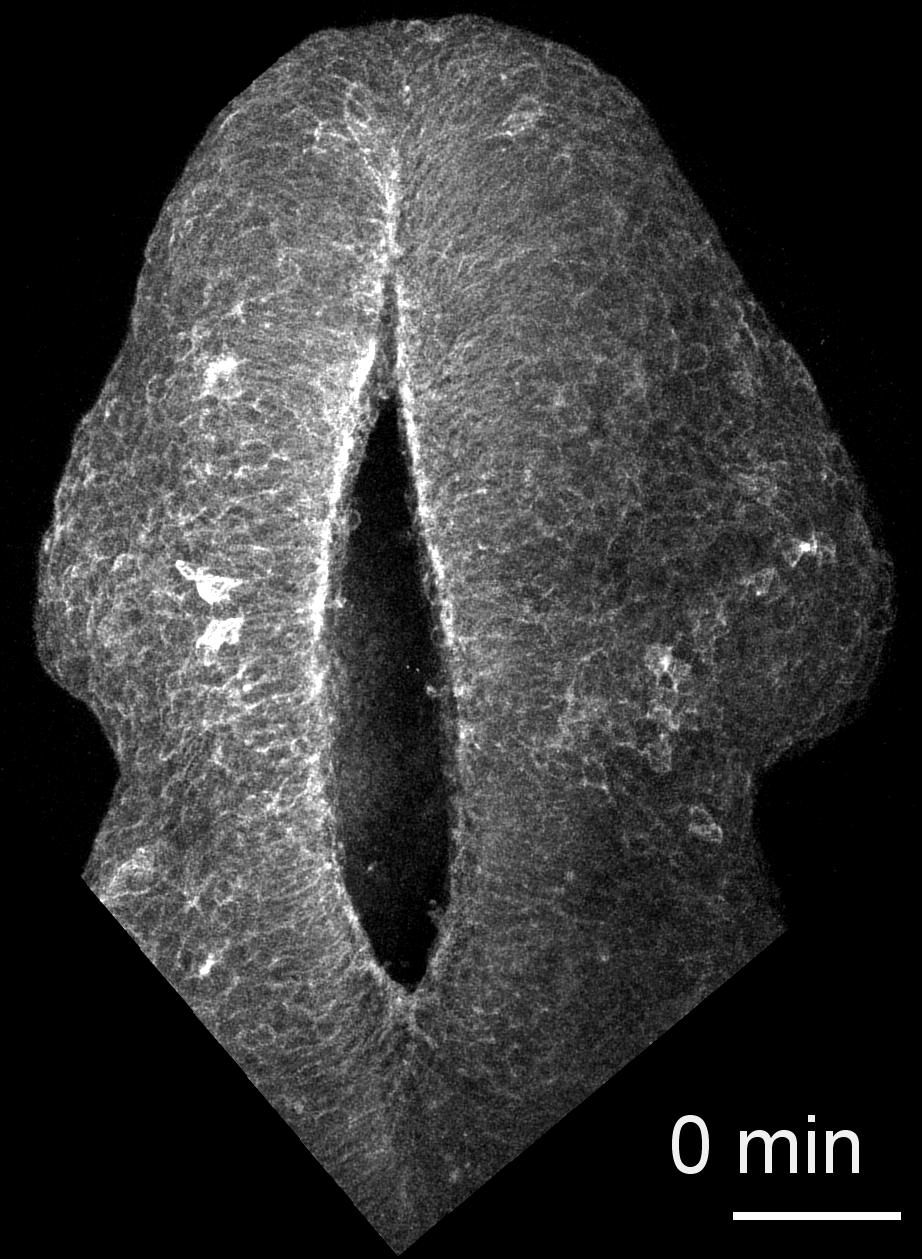

### Movie S3

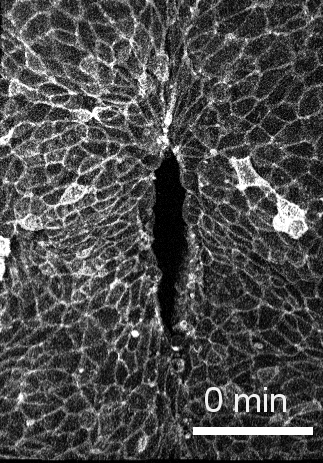

### Movie S4

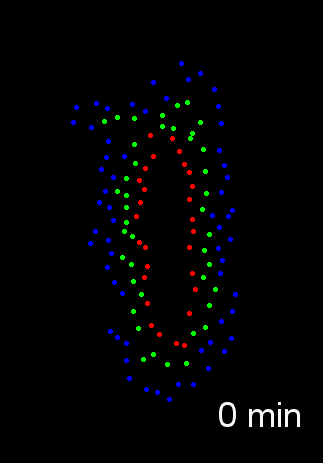
